# Contributions of the conserved insect carbon dioxide receptor subunits to odor detection

**DOI:** 10.1101/803270

**Authors:** Arun Kumar, Genevieve M. Tauxe, Sarah Perry, Christi A. Scott, Anupama Dahanukar, Anandasankar Ray

**Affiliations:** Department of Molecular, Cell and Systems Biology, University of California, Riverside Riverside, CA 92521, USA; Department of Entomology, University of California, Riverside Riverside, CA 92521, USA; Graduate Program in Genetics, Genomics, and Bioinformatics, University of California, Riverside Riverside, CA 92521, USA; Graduate Program in Cell, Molecular and Developmental Biology, University of California, Riverside Riverside, CA 92521, USA

**Author notes:** These authors contributed equally (Arun Kumar, Genevieve M. Tauxe).

## Abstract

The CO_2_ receptor in mosquitoes is broadly tuned to detect many diverse odorants. The receptor consists of three 7-TM subunits (Gr1, Gr2, and Gr3) in mosquitoes but only two subunits in *Drosophila*: Gr21a (Gr1 ortholog) and Gr63a (Gr3 ortholog). We demonstrate that *Gr21a* is required for CO_2_ responses in *Drosophila* as has been shown for *Gr63a*. Next, we generate a *Drosophila* double mutant for *Gr21a* and *Gr63a*, and in this background we functionally express combinations of *Aedes Gr1, 2*, and *3* genes in the “CO_2_ empty neuron.” Only two subunits, Gr2 and Gr3, suffice for response to CO_2_. Addition of Gr1 increases sensitivity to CO_2_ while it decreases the response to pyridine. The inhibitory effect of the antagonist isobutyric acid is observed upon addition of Gr1. Gr1 therefore increases the diversity of ligands of the receptor, and also modulates the response of the receptor complex.

## Introduction

Three members of the insect *Gustatory receptor* (*Gr*) family are expressed in olfactory neurons that detect airborne carbon dioxide (CO_2_) and other odorants: *Gr1, Gr2*, and *Gr3* (Jones et al., 2007; Kwon et al., 2007; Lu et al., 2007; MacWilliam et al., 2018; Tauxe et al., 2013; Turner et al., 2011; Turner and Ray, 2009). These receptors are conserved in holometabolous insect orders except Hymenoptera (MacWilliam et al., 2018; Robertson and Kent, 2009). In mosquitoes, they are expressed in specialized neurons called cpA, which detect CO_2_ (Jones et al., 2007; Lu et al., 2007; Syed and Leal, 2007) and a number of additional compounds, including many found in human skin odor (Lu et al., 2007; Tauxe et al., 2013; Turner et al., 2011; Turner and Ray, 2009). These neurons play a critical role in host seeking behavior of mosquitoes (Gillies, 1980; McMeniman et al., 2014; Tauxe et al., 2013; Turner et al., 2011). Apart from being an attractant itself, past studies have also established that just a transient exposure to a filamentous plume of CO_2_ “activates” behavior, and instantly lowers the threshold of response to human skin odor in *Ae. aegypti* by a factor of at least five (Dekker et al., 2005; Healy and Copland, 2000) and also increases attraction to heat (Liu and Vosshall, 2019; McMeniman et al., 2014) and visual cues (Vinauger et al., 2019).

Unlike all other insects studied thus far, *Drosophila* lack a Gr2 equivalent and instead form a functional CO_2_ receptor with Gr1 and Gr3 orthologs (Jones et al., 2007; Kwon et al., 2007; Robertson and Kent, 2009). Gr21a and Gr63a are co-expressed in ab1C neurons, one of four neurons housed in ab1 antennal basiconic sensilla (Figure 1A). When the fly CO_2_ receptor subunits are ectopically expressed in another neuron, both are required to confer CO_2_ responses (Jones et al., 2007), and ab1C neurons lacking *Gr63a* do not respond to CO_2_ or other odorants (Jones et al., 2007; Tauxe et al., 2013). To compare how these CO_2_ receptor subunits function across species, we generated mutant *Drosophila* that express combinations of mosquito Grs in the same neuron.

**Figure 1.**
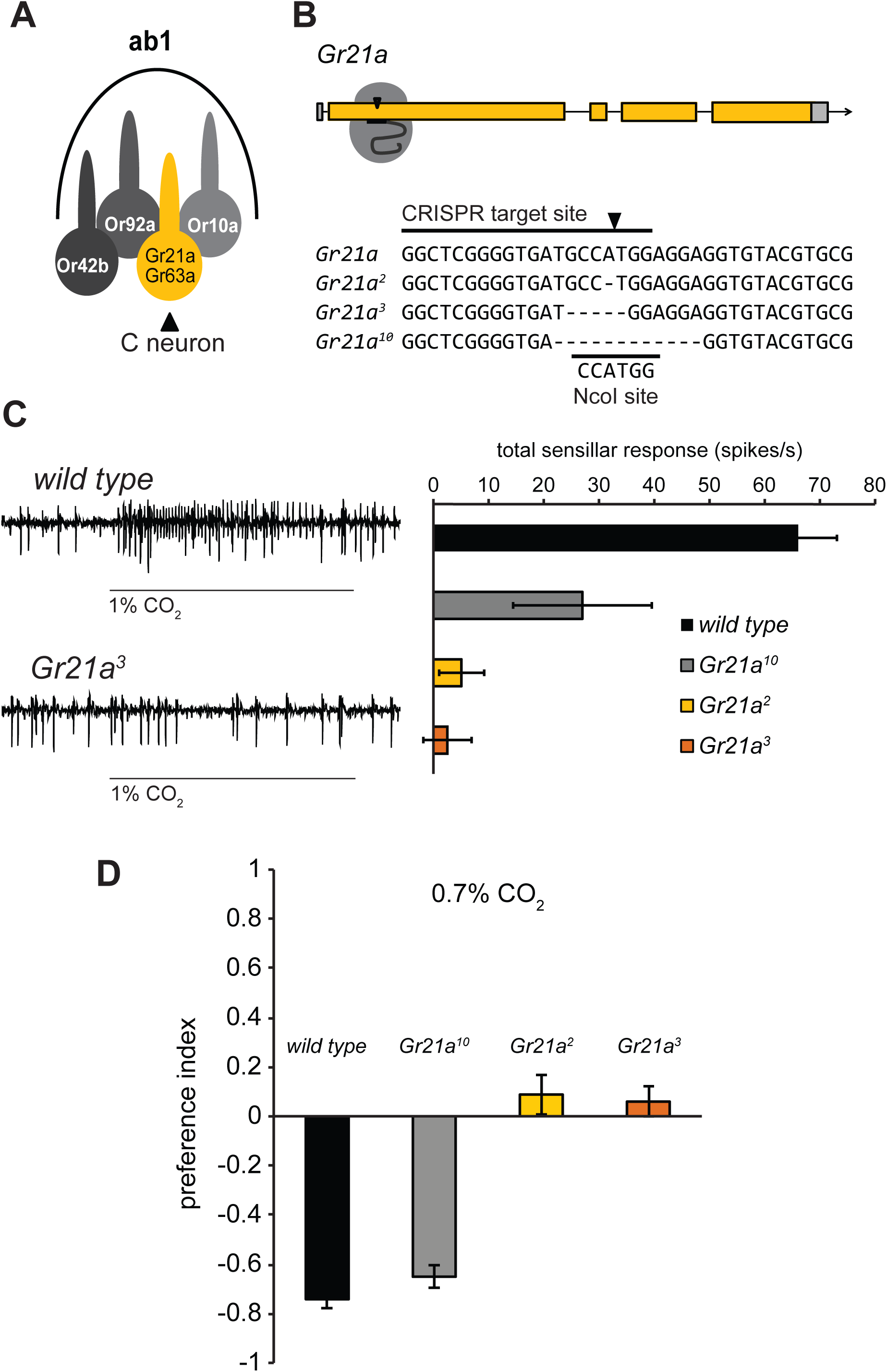
Gr21a is required for detection of CO_2_ in *Drosophila*. (**A)** The ab1 sensillum. (**B)** CRISPR target site in the first exon of *Gr21a* and sequences of three mutant alleles compared with the wild type *Gr21a* sequence from FlyBase. The NcoI restriction site is indicated. **(C)** Representative traces and mean ab1 sensillar responses to 1% CO_2_ in flies with different *Gr21a* alleles (*n* = 6 except *Gr21a*^10^, *n* = 3). **(D)** Behavioral responses to 0.7% CO_2_ (*n* = 10). **(C,D)** Error bars are s.e.m. See also Figure S1.

## Results

The requirement for *Gr21a* has not been directly tested via mutant analysis, so we used the CRISPR/Cas9 system to create small deletions early in the *Gr21a* coding region (Figure 1B). Out of 19 sequenced alleles, we selected three for analysis. *Gr21a*_*2*_ and *Gr21a*^*3*^ have frameshift mutations predicted to encode prematurely truncated peptides, and *Gr21a*^*10*^ has an in-frame deletion that removes codons for four amino acid residues near the N terminal. Electrophysiological responses to CO_2_ were eliminated in the ab1 sensillum of flies homozygous for the *Gr21a*_*2*_ or *Gr21a*^*3*^ alleles, but not *Gr21a*^*10*^ (Figure 1C). *Drosophila* normally avoid CO_2_ in T-maze tests. Consistent with the neuronal responses in the three mutants, *Gr21a*_*2*_ and *Gr21a*^*3*^ mutants no longer avoided CO_2,_ but *Gr21a*^10^ mutants still did (Figure 1D). All genotypes avoided the control repellent benzaldehyde (Figure S1).

The CO_2_ receptor subunits are well conserved, with amino acid sequence similarity across species comparable to broadly-expressed co-receptors of other chemoreceptor families (Robertson and Kent, 2009, Fig. 2A). Inspired by the empty neuron system used to decode Or receptors of mosquitoes using *Drosophila* (Carey et al., 2010; Hallem and Carlson, 2006), we combined the *Gr21a*^*3*^ mutant with *ΔGr63a* and *Gr63a–GAL4* lines to create a “CO_2_ empty neuron system” in ab1C (Figure 2B). Using single sensillum electrophysiology we measured the total spike activity of the ab1 sensillum and found that, as expected, a single *Aedes* Gr receptor did not confer response to CO_2_ (Figure S2). We then expressed combinations of *Ae. aegypti* receptors in the *ΔGr21a*;*ΔGr63a* double mutant background and found that only two combinations conferred CO_2_ responses: Gr2+Gr3 and Gr1+Gr2+Gr3. This held true both for 0.3% CO_2_ and for exhaled breath (Figure S2).

**Figure 2.**
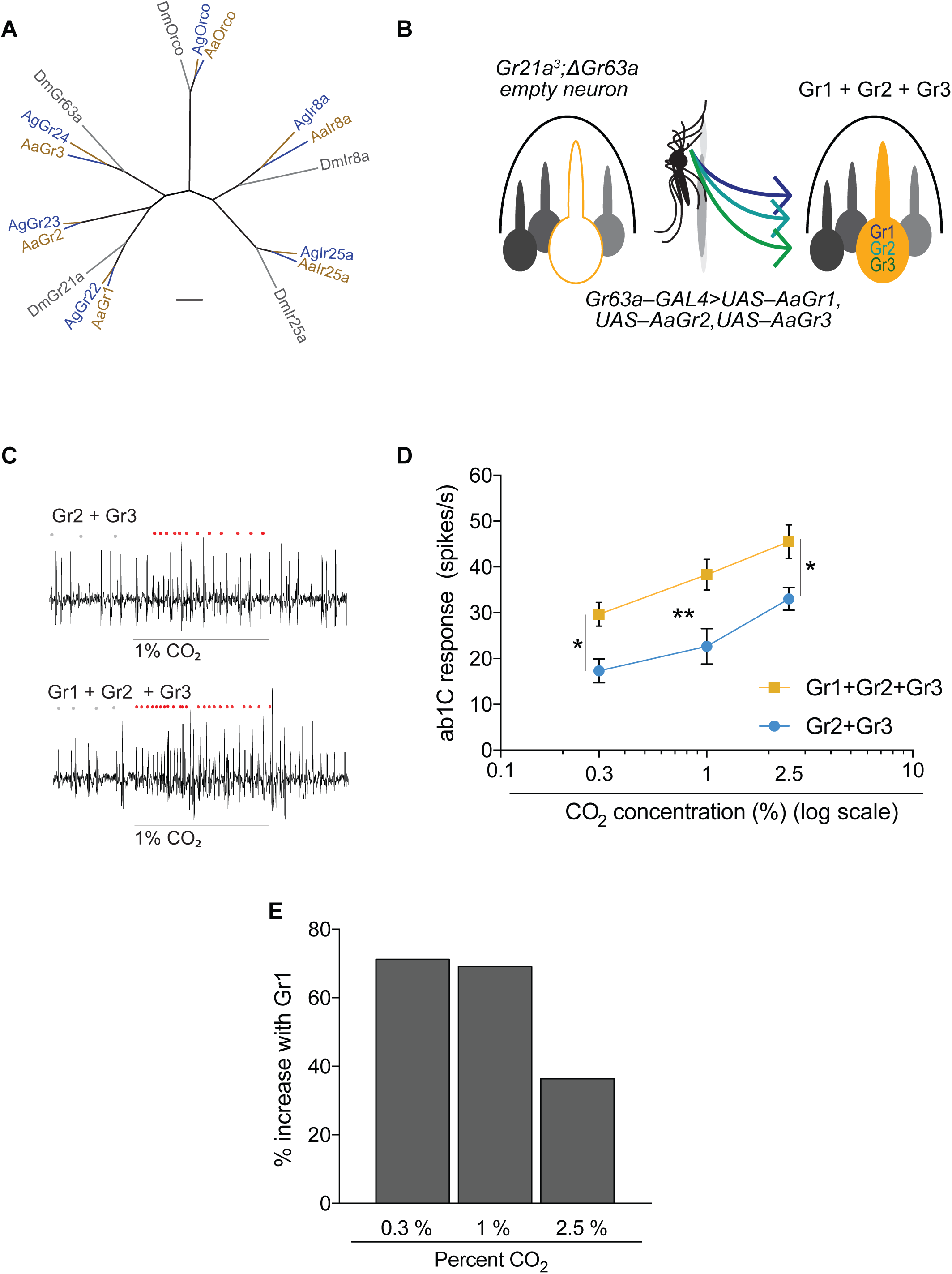
Gr1 increases CO_2_ sensitivity of the *Aedes* CO_2_ receptors. **(A)** Divergence of CO_2_ receptors and, for comparison, the odorant receptor co-receptor (Orco) and ionotropic receptor co-receptors Ir8a and Ir25a. Scale bar = 0.1 substitutions per unit length. **(B)** Schematic of CO_2_ empty neuron system. **(C)** Representative traces of the ab1 sensillum expressing mosquito receptors in the CO_2_ empty neuron with responses to a 0.5 s stimulus of 1% CO_2_. Dots mark action potentials attributed to the ab1C neuron during part of the baseline and the 0.5 s stimulus window. **(D)** Mean transgenic ab1C responses to increasing concentrations of CO_2_ (*n* = 6; 2-way ANOVA followed by Bonferroni multiple comparison test; \**p* < 0.05, \*\**p* < 0.01). Error bars are s.e.m. **(E)** Percent increase in CO_2_ response with the addition of Gr1 to Gr2+Gr3. See also Figure S2.

In order to more precisely measure odor-evoked responses, we singled out the activity of the C neuron in the ab1 sensillum in subsequent experiments. We found that Gr2+Gr3 was sufficient to restore CO_2_ sensitivity to the empty neuron in a dose-dependent manner (Figure 2 C,D). When we expressed all 3 *Aedes* Grs in the *Drosophila* CO_2_ empty neuron we observed a significant increase in response to CO_2_ across all concentrations tested (Figure 2C,D, Figure S3). Addition of Gr1 to Gr2+Gr3 increased the responsiveness of the neuron by ∼70% at the lower concentrations tested (Figure 2D,E).

The heteromeric nature of the receptor offers the potential to investigate the relative contributions of the different subunits to ligand detection in the ab1C expression system. Introducing a second copy of Gr1 (Gr1+Gr1+Gr2+Gr3) or doubling the copy number of Gr2+Gr3 in the presence of Gr1 (Gr1+Gr2+Gr2+Gr3+Gr3) increased the response of the ab1C neuron to 2.5% CO_2_ (Figure 3A). The simplest interpretation of these results is that Gr1 increases the sensitivity of the neuron to carbon dioxide.

**Figure 3.**
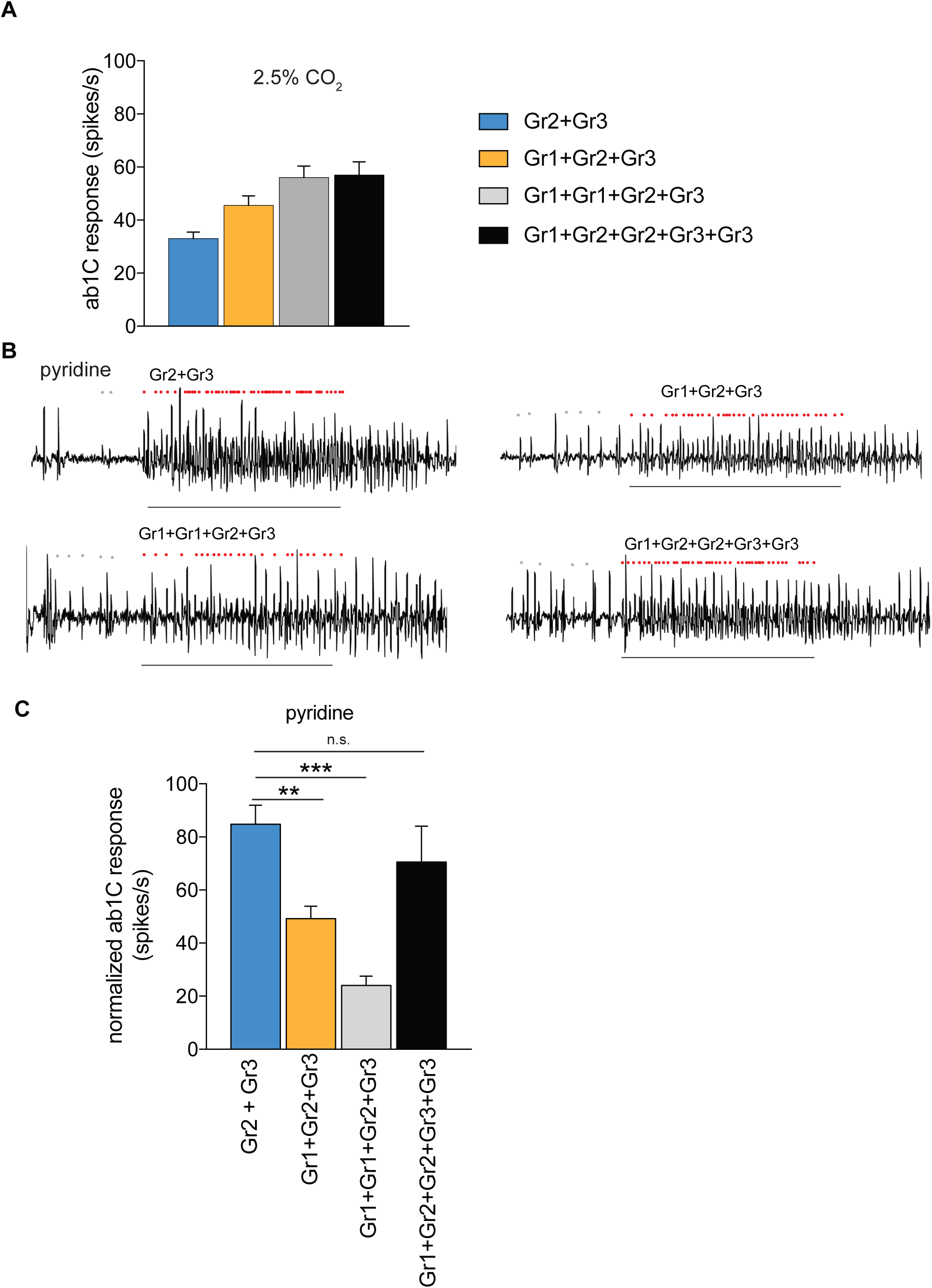
Odor responses of *Aedes aegypti* receptors in the CO_2_ empty neuron are modulated by Gr copy number. **(A)** Mean ab1C responses to 2.5% CO_2_ for different Gr combinations (*n* = 6–12). **(B)** Representative traces of ab1 sensilla expressing mosquito receptors in the CO_2_ empty neuron with responses to pyridine. Dots mark action potentials attributed to the ab1C neuron during part of the baseline and the 0.5 s stimulus window. **(C)** Mean odorant responses normalized to the total sensillar response in the CO_2_ empty neuron. Error bars are s.e.m. (*n* = 6–8; ANOVA followed by Dunnett’s test comparing results to Gr2+Gr3, \*\**p* < 0.01, \*\*\**p* < 0.001). **(B,C)** Pyridine was diluted at 1% (v/v) in paraffin oil. The copy number of each transgene is indicated; all used a single copy of the *Gr63a–GAL4* driver (see Table S1). Stimulus = 0.5 s. See also Figure S3.

In order to test whether copy number of Gr1 could increase sensitivity of the receptor complex to a non-CO_2_ agonist, we tested pyridine, which is an activator of the cpA neuron in mosquitoes. This odorant was selected from an initial screen of cpA agonists because we can count the activity of the C neuron without significant interference from the other 3 neurons in the ab1 sensillum. The expression of Gr2+Gr3 alone conferred a response to pyridine (Figure 3B,C). Contrary to what we found with CO_2_, the addition of Gr1 (Gr1+Gr2+Gr3) decreased the pyridine response significantly. Doubling the copy number of Gr1 further decreased the response to pyridine. The effect of Gr1 could be reversed by increasing the relative copy number of the other receptors (Gr1+Gr2+Gr2+Gr3+Gr3) (Fig 3B,C). These results indicate that Gr1 modulates the responses of the Gr2+Gr3 receptor in different ways depending upon the identity of the odorant: it increases sensitivity to CO_2_ but decreases the sensitivity to pyridine (Figure 3).

We also tested an inhibitory odorant, isobutyric acid, which inhibits CO_2_ responses in mosquito cpA neurons (Fig 4A,B). This odorant was also selected because we can count the activity of the C neuron in the ab1 sensillum. In order to test the effect of this inhibitor on responses of Grs expressed in the ab1C system, we overlaid a 2 s pulse of 0.7% CO_2_ with a 0.5 s pulse of the inhibitory odorant or water solvent alone. Isobutyric acid did not inhibit CO_2_ response in *Drosophila* ab1C neurons expressing only Gr2+Gr3 (Figure 4C,D). Interestingly, when Gr1 was co-expressed with Gr2+Gr3, isobutyric acid significantly inhibited the ab1C response to CO_2_ as compared to the solvent control (Figure 4C,D).

**Figure 4.**
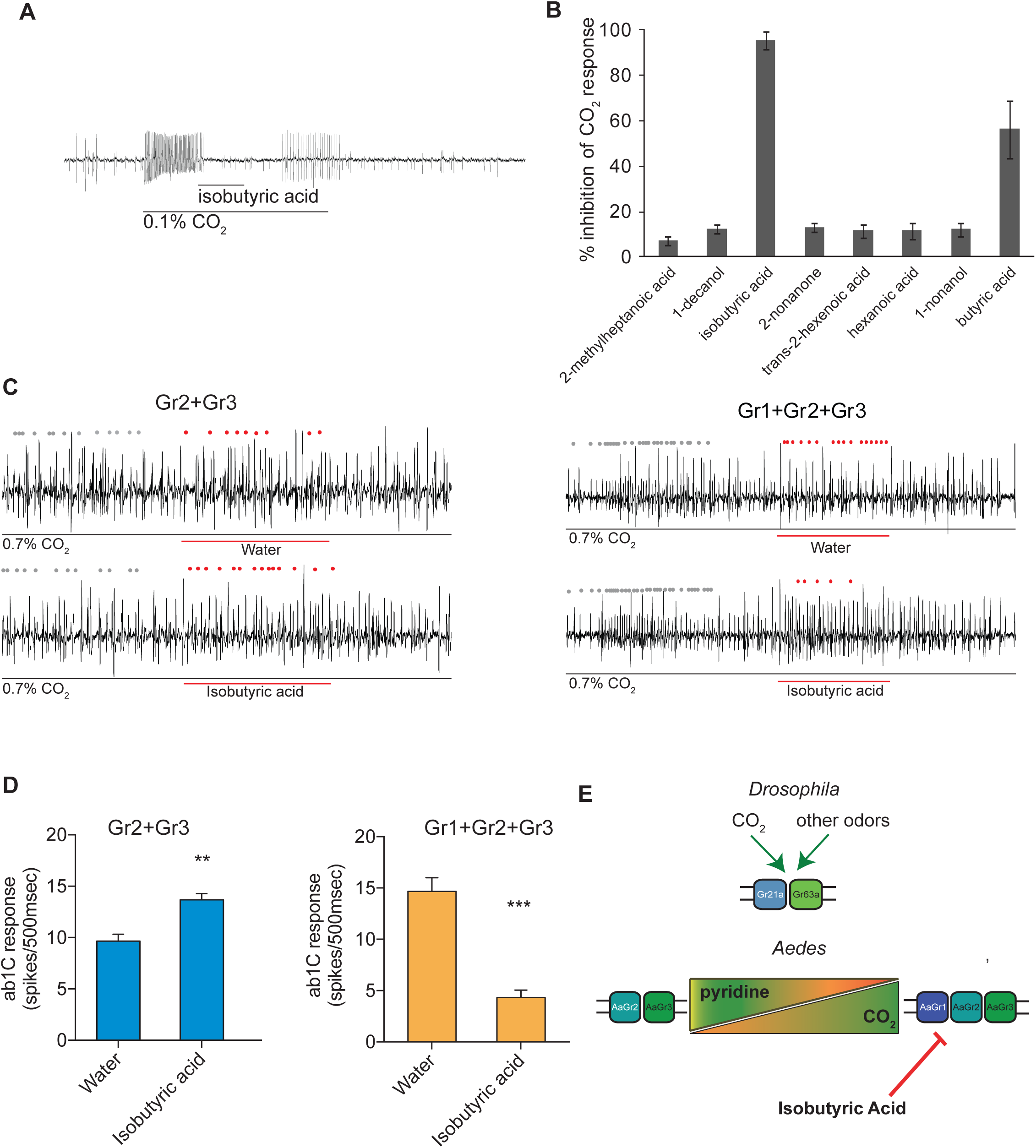
The *Aedes* CO_2_ receptor is inhibited by isobutyric acid only when Gr1 is present. **(A)** Representative traces of a cpA neuron in female *Aedes aegypti* responding to CO_2_ overlaid with isobutyric acid, and **(B)** mean percent inhibition of cpA responses by odorants relative to solvent. Odorants were diluted at 10% (v/v) in paraffin oil and presented as a 0.5 s stimulus during a 2 s pulse of 0.1% CO_2_ (*n* = 5–6). Error bars are s.e.m. **(C)** Representative traces of a CO_2_ empty neuron expressing mosquito Grs in *D. melanogaster* responding to a 2 s pulse of CO_2_ overlaid with 0.5 s of isobutyric acid or solvent. Dots mark action potentials attributed to the ab1C neuron during part of the baseline and the 0.5 s stimulus window. **(D)** Mean response of the transgenic ab1C neuron to inhibitor overlay in flies expressing combinations of mosquito Grs (*n* = 6–9; *t*-test, \*\**p* < 0.01, \*\*\**p* < 0.001). Odorants were diluted at 1% (v/v) in water and presented as a 0.5 s stimulus during a 2 s pulse of 0.7% CO_2_. Error bars are s.e.m. **(E)** Model of receptor subunit contributions to odorant response.

## Discussion

We conclude that the *Ae. aegypti* CO_2_ receptor can function as a heteromer of Gr2+Gr3, whose responses are modulated by Gr1. Responses to CO_2_ are increased and responses to pyridine are reduced, suggesting that activating odorants may be divided into two classes based on how Gr1 affects their detection. Additionally, CO_2_ responses are only effectively inhibited by isobutyric acid when AaGr1 is present. This is unlike the situation in wild type *Drosophila*, in which the ab1C neuron functions without a homolog of Gr2 (Robertson and Kent, 2009), suggesting that the two remaining *Drosophila* receptors have taken over some of AaGr2’s function. The potential functional differences between CO_2_ receptors of the fly and mosquito also stand in contrast with our knowledge of the odorant receptor (Or) family, in which the fly co-receptor Orco is compatible with ligand-specific Ors from multiple insect species, including mosquitoes (Carey et al., 2010; Hallem et al., 2004). A previous study showed that expressing the *An. gambiae* receptors AgGr22+AgGr24 (homologous to AaGr1+AaGr3) in *Drosophila* failed to elicit a response to 100% CO_2_ until either AgGr23 (homologous to AaGr2) was added or gene dosage of the other receptors was increased; however, unlike this study the ab3A neurons were used which normally express members of the Or gene family (Lu et al., 2007). Further experiments will be required to determine whether the difference in CO_2_ responses are due to a different neuronal environment, or a difference in receptor function between the two mosquito species.

We do not know as yet whether there are any mechanisms that regulate relative expression levels of the three Gr subunits of the CO_2_ receptor complex. However, our findings open the possibility that such mechanisms could modulate sensitivity to different agonists and antagonists, which in turn could alter behavior. Interestingly, expression analyses in *Anopheles* have revealed that mRNA levels of *AgGr22* (homolog of *AaGr1*), but not of the other two subunits (*AgGr23* and *24*), are substantially higher 4 days post-eclosion as compared to 1 day post-eclosion, which is correlated with a significant increase in sensitivity to CO_2_ over the same period of time (Omondi et al., 2015). The CO_2_ receptor subunits are among the most evolutionarily conserved of all insect chemosensory receptors, and they are activated or inhibited by a wide range of structurally diverse ligands (Tauxe et al., 2013; Turner et al., 2011, MacWilliam et al., 2018). Understanding the circumstances under which the responses of these receptors are modulated and the molecular mechanisms that are behind their functional versatility will add to our knowledge of basic mechanisms of olfactory receptor function and aid in the design of new tools to manipulate the host-seeking behavior of mosquitoes and other insects.

## Experimental Procedures

### Insects

*Drosophila melanogaster* were reared on standard cornmeal–dextrose medium at 25°C. Receptors were cloned from *Aedes aegypti* Orlando strain mosquitoes.

### Generation of mutant and transgenic fly lines

The CRISPR target site was chosen using tools available online at http://flycrispr.molbio.wisc.edu/tools. Targeting oligos CTTCGGCTCGGGGTGATGCCATGG and AAACCCATGGCATCACCCCGAGCC were ligated directly into BbsI-digested pU6-BbsI-chiRNA (Addgene #45946). Resulting clones were screened for addition of a NcoI restriction site. The U6-Gr21a-chiRNA cassette was removed using KpnI and EcoRI digestion and cloned into similarly cut pattB. The resulting pattB{U6-Gr21a-chiRNA} vector was used to transform y,w;attP40 embryos using the site-directed phiC31 integrase system. Vas-Cas9 (BL# 51324) females were mated with attP40{Gr21a-chiRNA} males and the resulting attP40{Gr21a-chiRNA}/+; vas-Cas9/+ females were mated to balancer males to generate 19 isogenic lines. 19/19 lines exhibited indels at the CRISPR target site: 16 frame-shift alleles and 3 in-frame deletion alleles.

Messenger RNA was isolated from the mouthparts of 5-day-old adult, non-bloodfed female *Ae. aegypti*. Gr genes were amplified from reverse transcribed cDNA by PCR using proof-reading enzymes, cloned and sequenced. Two isoforms of AaGr2 were present in the source cDNA, which differ in length by 75 nucleotides at the 5′ end of the coding sequence; the shorter allele is expressed with greater frequency and was cloned using a modified pENTR system (“pATTL”) developed by G. M. Pask. The gene fragments were inserted into pUASg-attB-DV vectors, which were used to transform flies using the site-directed phiC31 integrase system. Presence of the UAS–AaGr construct in the resulting transgenic lines was confirmed by PCR. Injections for both sets of flies were carried out by Genetic Services, Inc. (Sudbury, MA). Complete genotypes and sources of flies used are listed in Supplemental Experimental Procedures.

### Electrophysiology

Single-sensillum recordings were performed essentially as described (Tauxe et al., 2013). Adult female *Drosophila* were tested 4–8 days after emergence. Briefly, insects were restrained and saline-filled glass micropipet electrodes were inserted into the test sensillum (recording electrode) and the compound eye (reference electrode). Signals were amplified 1000×, band-pass filtered, and digitized to record action potentials of the neurons in the test sensillum. The identity of ab1 sensilla was confirmed at the beginning of all recordings by presence of more than 2 neuronal spike amplitudes as well as by shape and location of sensilla. Up to 3 sensilla were tested per insect.

Chemicals were obtained from Sigma-Aldrich at the highest available purity, typically >98%. For each odorant stimulus odor cartridges were made with a Pasteur pipette attached to a 1ml pipet tip, and 50µl of the odorant solutions in indicated concentrations and solvents was added to a cotton wool piece. Stimuli were presented in roughly the same order across replicates; each cartridge was used for ≤3 stimuli. A constant 5 ml/s stream of carbon filtered room air was switched from a blank cartridge to the odor cartridge using a Syntech CS-55 to present odor stimuli. The resulting airflow was delivered into a glass tube with a constant, humidified airstream (10 ml/s) whose mouth was centered on and ∼1 cm from the insect head. CO_2_ stimuli were pulsed using a PM8000 microinjector (MicroData Intrument, Inc.) or MNJ-D microinjector (Tritech Research) to deliver controlled pulses from pressurized cylinders of 1% and 5% CO_2_ in air into the carrier airstream, resulting in the indicated final concentration of gas at the insect head.

In the ab1 sensillum of *D. melanogaster* and the cp sensillum of *Ae. aegypti*, isolated action potentials from individual neurons can be distinguished by their relative magnitude. In all agonist experiments, to correct for baseline firing rate, sensillar responses were calculated as 2 × (number of spikes during 0.5 s stimulus presentation) – (number of spikes during 1 s prior to stimulus presentation). In the preliminary survey shown in Figure S2, exhaled breath stimuli were provided by the experimenter blowing directly on the fly with other airflows held constant. VU0183254, also known as VUAA-ANT, is a specific antagonist of Orco (Jones et al., 2012) and was present in the recording electrode saline during some of these recordings. It had no effect on the magnitude of odorant responses (two-way ANOVA with type III sum of squares; main effect and interaction of VU0183254 *p* > 0.05 for all test odorants). Data with and without this compound have been pooled in Figure S2.

In subsequent experiments, individual *D. melanogaster* ab1C spikes and *Ae. aegypti* cpA spikes were counted and corrected for baseline activity as above. For pyridine recordings, the total activity of the ab1 sensillum was counted, and the activity attributable to the ab1C neuron determined by subtracting the activity of the ab1 sensillum in the *Gr21a*^*3*^/*ΔGr63a* double mutant.

For inhibition experiments with isobutyric acid, a 2-s CO_2_ pulse was overlaid with a 0.5 s pulse of 1% isobutyric acid or solvent alone (water). The activity of ab1C was counted for 500 ms in both, as described previously (Turner and Ray, 2009). Percent inhibition was calculated relative to the response to the solvent control in the same sensillum.

Since *Ae. aegypti* Gr constructs were inserted into the *Drosophila* genome at two different locations, data presented in Figures 3,4, S2, S3 for genotypes with three or more UAS constructs include multiple positional variants. In initial screens, gene position had no effect on ab1C odor responses (Figure S2; two-way ANOVA of responses across odors and responding genotypes with three UAS constructs; main effect and interaction of gene position *p* > 0.05), so these data were pooled for analysis.

### *Drosophila* behavior

T-maze assays were performed essentially as described (Turner and Ray, 2009). For each trial, 40 flies (20 males and 20 females, 3–7 days old) were starved with access to water for 24 hrs, loaded into an apparatus, and allowed to choose freely between two arms of the maze: 15 ml culture tubes, one with an odor stimulus and one without, for 1 min. No air flow was used. For CO_2_ stimuli, 100 μl pure CO_2_ was injected through the cap of the stimulus tube via syringe immediately before attaching the arms to the maze for each trial; reported concentration is an approximation based on injected volume. Ambient room CO_2_ was present in both arms. For benzaldehyde stimuli, 10 μl of 10% benzaldehyde in paraffin oil was applied to a small disc of filter paper inserted into the test arm of the T-maze immediately before each trial; an equivalent disc with 10 μl paraffin oil was inserted into the control arm. Preference index is calculated as: (number of flies in test arm – number of flies in control arm) / (total number of flies in both arms) and ranges from 1 (perfect attraction) to -1 (perfect avoidance).

### Genetic Analysis

The phylogenetic tree in Figure 2A was generated in ClustalW 2.0 (Larkin et al., 2007) from amino acid sequences.

## Acknowledgements

We thank Erica Freeman for assisting in electrophysiology recordings. We thank Gregory Pask who designed a cloning vector. We thank support from NIAID (NIH) for this project, award R01AI087785 to A.R.

## SUPPLEMENTAL FIGURE LEGENDS

**Figure S1, related to Figure 1.** Behavioral responses to benzaldehyde in *Gr21a* mutants (*n* = 10 trials). Error bars are s.e.m.

**Figure S2, related to Figure 2**

Schematic, representative traces, and mean responses of ab1 sensilla expressing mosquito receptors in the CO_2_ empty neuron to 0.5 s stimuli of CO_2_ or puffs of exhaled breath. One copy of each indicated transgene was present. Dots mark action potentials attributed to the ab1C neuron. (*n* = 6–28; ANOVA followed by Dunnett’s test comparing results to empty neuron control, \**p* < 0.05, \*\**p* < 0.01, \*\*\**p* < 0.001). Error bars are s.e.m.

**Figure S3, related to Figure 3**

**(A)**Representative traces of the ab1 sensillum expressing mosquito receptors in the CO_2_ empty neuron with responses to a 0.5 s stimulus of 1.67% CO_2_. Dots mark action potentials attributed to the ab1C neuron during part of the baseline and the 0.5 s stimulus window. **(B)** Mean transgenic ab1C responses to increasing concentrations of CO_2_ (*n* = 8). **(C)** Representative traces of the baseline response of ab1 sensillum expressing mosquito receptors in the CO_2_ empty neuron. **(D)** Mean ab1C baseline activity in spikes/s.

