## Supplemental Table 1 for "Contributions of the conserved insect carbon dioxide receptor subunits to odor detection"

**Table S1. Full genotypes and sources of flies used in transgenic experiments.**

Alleles used in transgenic experiments.

| name | full allele designation | notes |
|---|---|---|
| wild type | | wCS (white Canton S) background |
| AaGr1 | *UAS–AaGr1A16* | ΦC31 injection, attP40 site (2nd chromosome) |
| AaGr1 | *UAS–AaGr1C49* | ΦC31 injection, VK00027 site (3rd chromosome) |
| AaGr2 | *UAS–AaGr2A10* | ΦC31 injection, attP40 site (2nd chromosome) |
| AaGr2 | *UAS–AaGr2C45* | ΦC31 injection, VK00027 site (3rd chromosome) |
| AaGr3 | *UAS–AaGr3A2* | ΦC31 injection, attP40 site (2nd chromosome) |
| AaGr3 | *UAS–AaGr3C46* | ΦC31 injection, VK00027 site (3rd chromosome) |
| ΔGr21a | $Gr21a^3$ | CRISPR deletion, see Fig. 1 |
| ΔGr63a | *ΔGr63a* | Bloomington *Drosophila* Stock Center 9941 |
| GAL4 | *Gr63a–GAL4* | Bloomington *Drosophila* Stock Center 9942 |

Full genotypes of transgenic flies. Allele/construct names are as above.

| fly name | full genotype |
|---|---|
| empty neuron | *w; ΔGr21a; ΔGr63a* |
| DmGr21a + AaGr3 | *w; AaGr3; ΔGr63a,GAL4* |
| AaGr1 + DmGr63a | *w; ΔGr21a,AaGr1; GAL4* |
| AaGr2 + DmGr63a | *w; ΔGr21a,AaGr2; GAL4* |
| AaGr1 + AaGr2 + DmGr63a | *w; ΔGr21a,AaGr1/ΔGr21a,AaGr2; GAL4* |
| *UAS–AaGr1,UAS–AaGr2,UAS–AaGr3* | *w; ΔGr21a,AaGr1/ΔGr21a,AaGr2; ΔGr63a,AaGr3/ΔGr63a*<br>*w; ΔGr21a,AaGr1/ΔGr21a,AaGr3; ΔGr63a,AaGr2/ΔGr63a*<br>*w; ΔGr21a,AaGr2/ΔGr21a,AaGr3; ΔGr63a,AaGr1/ΔGr63a* |
| *Gr63a–GAL4* | *w; ΔGr21a; ΔGr63a/ΔGr63a,GAL4* |
| Gr1 | *w; ΔGr21a,AaGr1/ΔGr21a; ΔGr63a/ΔGr63a,GAL4* |
| Gr2 | *w; ΔGr21a,AaGr2/ΔGr21a; ΔGr63a/ΔGr63a,GAL4* |
| Gr3 | *w; ΔGr21a,AaGr3/ΔGr21a; ΔGr63a/ΔGr63a,GAL4* |
| Gr1 + Gr2 | *w; ΔGr21a,AaGr1/ΔGr21a,AaGr2; ΔGr63a/ΔGr63a,GAL4* |
| Gr1 + Gr3 | *w; ΔGr21a,AaGr1/ΔGr21a,AaGr3; ΔGr63a/ΔGr63a,GAL4* |
| Gr2 + Gr3 | *w; ΔGr21a,AaGr2/ΔGr21a,AaGr3; ΔGr63a/ΔGr63a,GAL4*<br>*w; ΔGr21a,AaGr2/ SΔGr21a; ΔGr63a,AaGr3/ΔGr63a,GAL4* |
| Gr1 + Gr2 + Gr3 | *w; ΔGr21a,AaGr1/ΔGr21a,AaGr2; ΔGr63a,AaGr3/ΔGr63a,GAL4*<br>*w; ΔGr21a,AaGr1/ΔGr21a,AaGr3; ΔGr63a,AaGr2/ΔGr63a,GAL4*<br>*w; ΔGr21a,AaGr2/ΔGr21a,AaGr3; ΔGr63a,AaGr1/ΔGr63a,GAL4* |

| **fly name** | **full genotype** |
|---|---|
| Gr2 + Gr2 + Gr3 + Gr3 | *w; ΔGr21a,AaGr2/ ΔGr21a,AaGr2; ΔGr63a,AaGr3,GAL4/ΔGr63a,AaGr3* |
| Gr1 + Gr1 + Gr2+ Gr3 | *w; ΔGr21a,AaGr1/ ΔGr21a,AaGr1; ΔGr63a,AaGr3,GAL4/ΔGr63a,AaGr2* |
| Gr1 + Gr2 + Gr2 + Gr3 + Gr3 | *w; ΔGr21a,AaGr3/ ΔGr21a,AaGr2; ΔGr63a,AaGr1,AaGr2/ΔGr63a,AaGr3,GAL4* |
