## Supplemental Figure 1 for "Contributions of the conserved insect carbon dioxide receptor subunits to odor detection"

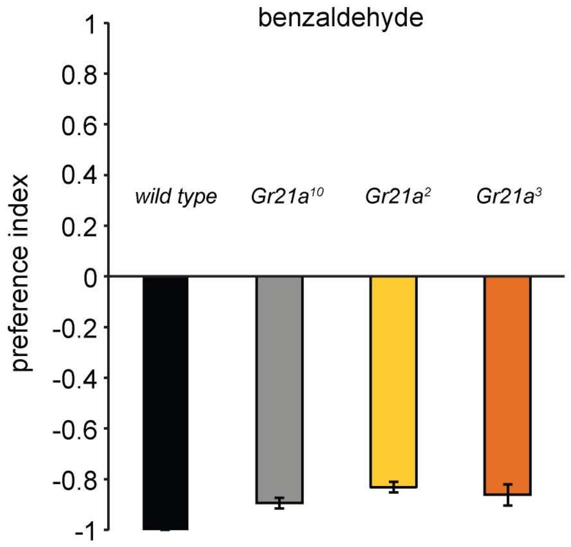


**Figure S1. Related to Figure 1.**
Behavioral responses to benzaldehyde in Gr21a mutants (n = 10). Error bars are s.e.m.
