## Supplemental Figure 2 for "Contributions of the conserved insect carbon dioxide receptor subunits to odor detection"

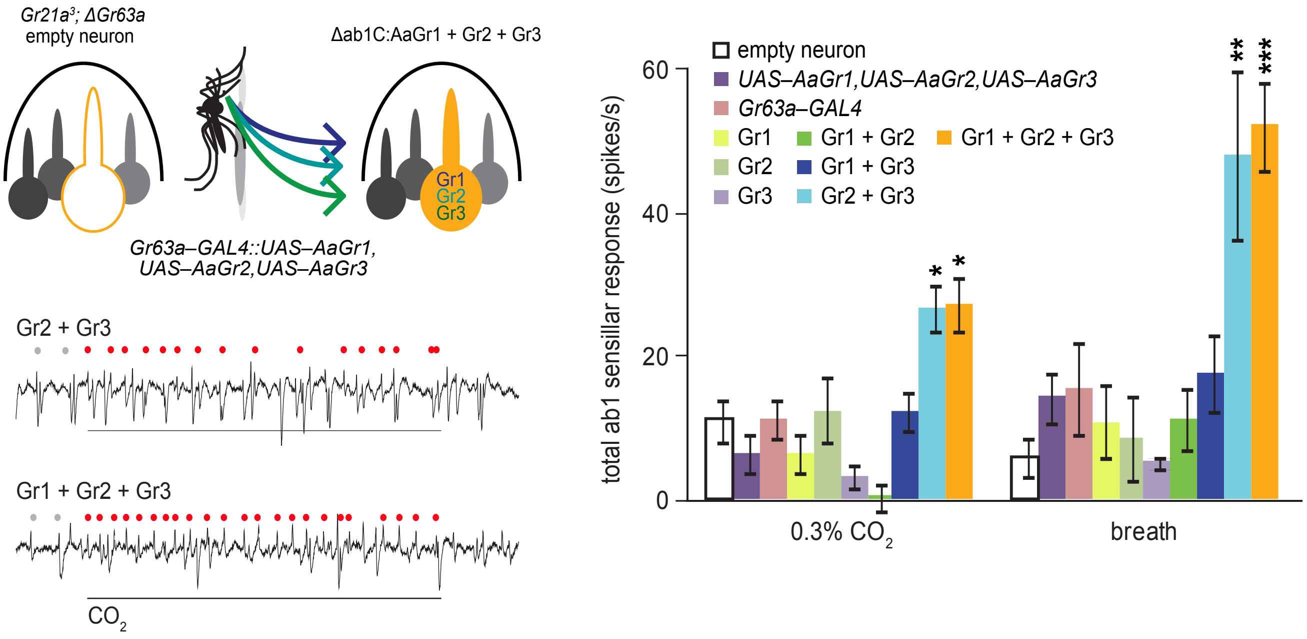


**Figure S2. Related to Figure 3.**

Schematic, representative traces, and mean responses of ab1 sensilla expressing mosquito receptors in the CO2 empty neuron to 0.5 s stimuli of CO2 or puffs of exhaled breath. One copy of each indicated transgene was present. Dots mark action potentials attributed to the ab1C neuron. (n = 6–28; ANOVA followed by Dunnett's test comparing results to empty neuron control,
*p < 0.05, **p < 0.01, ***p < 0.001). Error bars are s.e.m.
