## Supplemental Figure 3 for "Contributions of the conserved insect carbon dioxide receptor subunits to odor detection"

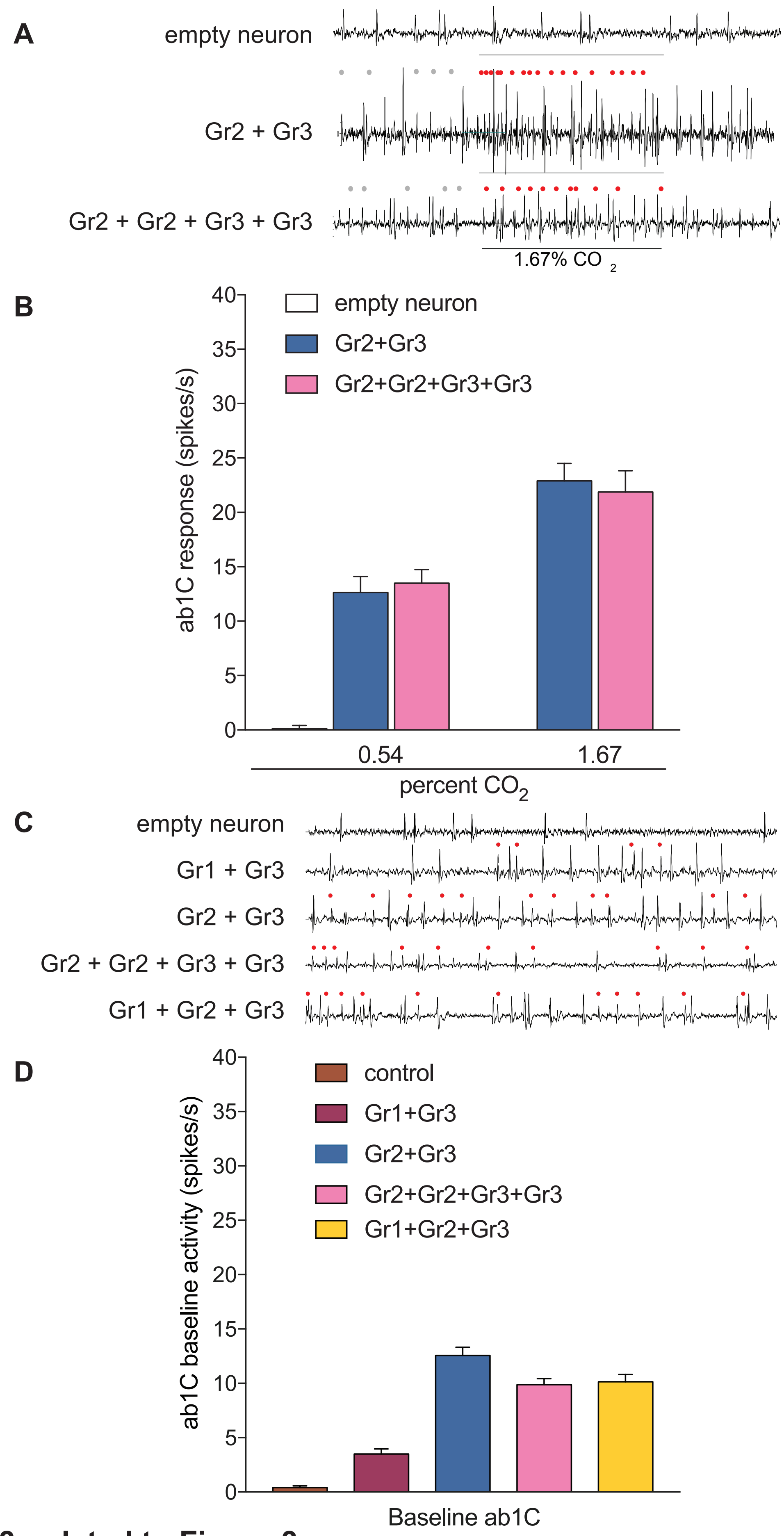


### Figure S3, related to Figure 3

**(A)** Representative traces of the ab1 sensillum expressing mosquito receptors in the $CO_2$ empty neuron with responses to a 0.5 s stimulus of 1.67% $CO_2$. Dots mark action potentials attributed to the ab1C neuron during part of the baseline and the 0.5 s stimulus window. **(B)** Mean transgenic ab1C responses to increasing concentrations of $CO_2$ (n = 8). **(C)** Representative traces of the baseline response of ab1 sensillum expressing mosquito receptors in the $CO_2$ empty neuron. **(D)** Mean ab1C baseline activity in spikes/s.
